# Biotic Assessment of Crowdsourced Data Defines Four Ecoregions in Thar: A Novel Approach for Citizen Engagement in Conservation

**DOI:** 10.1101/2023.04.13.536827

**Authors:** Manasi Mukherjee, Angshuman Paul, Mitali Mukerji

## Abstract

Distinct ecosystems have representative biota that can allow delineation of ecoregions. Despite being one of the most populated desert ecosystems in the world with extreme climatic conditions, Thar has been classified as a single ecoregion by WWF. The present study aimed to test this on the basis of biotic assessment using a citizen science-based crowdsourced data from ebird (492 species of birds through 50,000 checklists of 4000 birders). Unsupervised clustering (*k*=4) and mathematical validation using cosine similarity revealed the presence of four distinct ecoregions- Eastern Thar (ET), Western Thar (WT), Transitional Zone (TZ) and Cultivated Zone (CZ) in Thar. Most strikingly, these ecoregions when overlaid on geographical regions of Thar show that the CZ ecoregion was split between three distant and distinct geographic regions. Further spatial diversity estimates show CZ had the least *α* diversity (273) and highest *β* diversity (1.8) indicating the least similarity between the districts that comprised this ecoregion. The results suggest CZ as an evolving ecoregion or a consequence of habitat fragmentation due to anthropogenic effects. The presence of least number of exclusive species in CZ (0.2%) and TZ (0.8%) including the near-threatened species like *Laticilla burnesii* (Rufous-vented grass babbler) and *Pelecanus philippensis* (Spot-billed pelican) and an endangered species *Athene blewitti* (Forest Owlet) highlights the need for restoration of these threatened biota and ecoregions. This study for the first time proposes crowd-sourced bird data as an important biota-based tool to understand (a) established ecoregions in an ecosystem, (b) anthropogenic effects of agri-farmlands and (c) relation between geographic regions and biota.

## 1. Introduction

Residential biological organisms are a part of an essential tool-kit for biotic assessment of ecosystems. This allows evaluation of the potential effects of actions on species and their critical habitats. By recognizing and explicitly characterizing spatial diversity of species in ecosystems, it is possible to define and better explain distinct ecoregions(Maestre et al., 2021). Though propitious, biotic assessments also demand wise choice of representative taxa and their extensive spatio-temporal data. Amongst all, birds are considered as the most reliable environmental indicators (Smits and Fernie, 2013). Serving as flagships for habitats and ecosystems under threat, birds act as early warning systems for risk factors (Lovette and Fitzpatrick, 2016; Kovařík et al., 2021). Ecological information of birds have also become the most abundantly available crowd-sourced data globally because of motivated citizens (eBird, 2023). Harvesting this data meaningfully can help assess anthropogenic activities, especially in vulnerable ecoregions for targeted management and restoration. Desert ecosystems or arid lands being most vulnerable to climate change, (Sala et al., 2000) deserve special attention in biodiversity conservation. The great Indian desert, Thar covers an area of about 385,000 sq km (9% of India’s land area)(ul Islam and Rahmani, 2011) and is one of the most populated deserts of the world. Mostly seen as a barren wasteland, Thar is biologically a rich habitat, inhabited by 682 species of flora, and 1195 (2.12 % of Indian fauna) species of fauna. Though there are a number of protected areas like Desert National Park (DNP), Gajner Wildlife Sanctuary and Tal Chhapar Wildlife Sanctuary, there is an increased risk of habitat loss for its biota. Invasive species, anthropogenic activities, increased agri-farmlands, rapid urbanization, unplanned infrastructure, pollution, climate change, etc., are threats to its diversity (Bharadwaj and rafi Rahmani, 2020) and abundance of original species (Roy and Roy, 2019) Some of the critically endangered species of Thar like the Great Indian Bustard are rapidly dwindling in numbers and have faced fragmented population, possibly due to cumulative effect of anthropogenic effects and climate change. Though geographically diverse and distinct with respect to natural communities, this has been defined as ‘one’ ecoregion by Olson et al. (2001). Of the 867 terrestrial ecoregions listed by World Wildlife Fund (WWF), Thar is designated as an ecoregion of Deserts and xeric shrublands biome within Indomalayan realm. Surprisingly, Thar Desert does not feature amongst the 238 prioritized ecoregions for global conservation, that include six major deserts of the world along with central Asian deserts (Olson et al., 2001). Not only is Thar geographically and biologically diverse, but it is also a habitat for a number of critically endangered and near-extinct species. Though a part of Thar namely Rann of Kutch flooded grasslands have been included under the critically endangered flooded grassland and savannas category (Global 200 status), Thar in its entirety has no representation. This begs the question whether Thar is a comparatively more stable desert or is it an outcome of an unassessed and understudied desert ecosystem?

A major reason for the dearth of ecological studies in such a diverse ecosystem is the requirement of large-scale and diverse ecological data. Often facing constraints of time, resources, and expertise, such research has sought solutions from citizen Science and crowd sourced data. Though citizen science has gained pace in many scientific domains, from astronomy to medicine (Lewandowski et al., 2017), it has been majorly acknowledged in ecological and environmental sciences. Citizen science based crowd sourced data has been used for a wide range of ecological research viz biodiversity assessment, ecosystem health monitoring and pollution analysis (Fraisl et al., 2022). ebird.org by the Cornell lab of ornithology is one of the largest global biological citizen science program for bird diversity. Due to open access to high volume, regularly updated and broad spatial coverage data on bird taxa, ecologists have widely used ebird data for both quantitative and qualitative ecological analysis (Sullivan et al., 2014). With an aim to identify and characterize ecoregions in Thar desert, the present study has curated bird diversity data of 33 districts of Rajasthan from ebird.org.

We hypothesised that given the geographical diversity of the Thar, there would be multiple ecoregions that could be reflected through their biotic components and birds could be ideal ecological indicators for this assessment. To address this, we have used the crowd sourced data from ebird from 33 districts of Rajasthan(70% of Thar)(eBird, 2023). This data comprises of 50,000 checklists of more than 4000 birders. Unsupervised clustering of the data indicated the presence of four ecoregions in Thar that was also validated through cosine similarity matrix. Further, the *α, β* and *γ* diversity using sets revealed the most exclusive species of each delineated ecoregions. Surprisingly, one of the delineated ecoregions was formed between 6 districts that were geographically distant as well as distinct. This suggests, presence of similar bird diversity between these geographically distant and distinct districts could be due to emerging habitats as a consequence of altered crop patterns. This is a first ever study on single taxa based biotic-ecoregion delineation of a desert ecosystem.

## 2. Methods

### 2.1. Study area

In India, the major fraction (70%) of Thar desert is in Rajasthan, followed by the states of Gujarat, Punjab and Haryana (15%). The present study was conducted across 33 districts of Rajasthan spanning a geographical area from 27.4695° N to 70.6217° E (Fig. 1). Aravallis, the oldest mountain range of India demarcates Thar geographically into a Western (WT) and Eastern Thar (ET). The north-west region covered by blankets of blown sand is dry and arid with scarce vegetation. The north-east is comparatively fertile and habitable land forming the wet Thar (Choudhary, 2021). Eastern Thar is also characterized by wide variety of metamorphic schists, quartzites, marbles, gneiss and granite rocks. There is a transitional zone that comprise an area between Aravallis and WT on one side and ET on the other (Fig. 3b) and governs variability in rainfall. Due to this, the density of vegetation increases from west to east in scattered clumps or patches. The cultivated zone (CZ) comprises of six districts located in two distant regions. One part of the CZ encompasses the districts of Dungarpur, Jhalawar, Baran and Banswara located within the Chambal ravine region towards south-east of Rajasthan. This region is characterized by stony uplands and hillocks dominated with black soil. The other part spans Hanumangarh and Sri Ganganagar located in the Canal region (Sharma and Mishra, 2021), north of Rajasthan, within Ghaggar plains (Misra, 1967). This is a combined flat plain of two major rivers Ghaggar and Harka, dominated by alluvial soil. On the basis of Agro-Climatic Zones of Rajasthan (Misra, 1967), Hanumangarh and Sri Ganganagar are categorized as Irrigated North-western plain, Banaswara and Dungarpur under Humid southern plains and Baran as well as Jhalawar under Humid southeastern plains.

**Figure 1:**
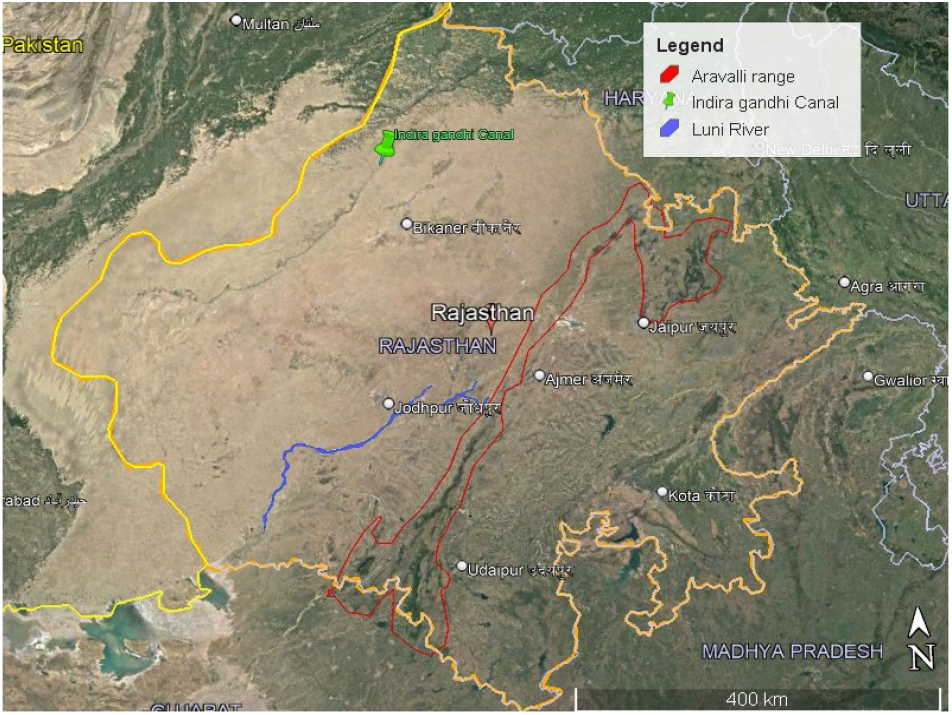
Satellite map of Thar Desert, Rajasthan (Source: Google Earth Pro, 2022). The Aravalii Range indicated with red line demarcates the sandy brown part of Western Thar (WT) from the comparatively greener Eastern Thar (ET).The blue line indicates the Luni river, the only river flowing through the dry and arid WT. The Indira Gandhi canal indicated with a green placemark, runs through the northwestern of Rajasthan’s Thar.

### 2.2. Data collection

Avian diversity data from 33 districts of Rajasthan were obtained from ebird.org, managed by the Cornell Lab of Ornithology. The website hosts crowd sourced data from birders around the world. eBird uses semistructured community science techniques with established methodologies for accounting for bias and effort within the data (Strimas-Mackey et al., 2020). The bird diversity data of Rajasthan in ebird.org comprising around 50,000 checklists by more than 4000 birders was curated for this study.

The data was curated from ebird.org in a matrix of columns with 492 species of birds in 33 districts of Thar desert, Rajasthan. We considered only the occurrence (presence or absence) data for species range and ecoregion delineation studies within Thar. Due to gaps in available count data, species abundance was not considered..

### 2.3. Analysis

The data was arranged to reflect the presence and absence of different bird species in different districts. We placed the bird species in columns and the districts in rows. For each row-column entry, if the corresponding bird species was found in a district, we indicated that by putting a ‘1’. Otherwise, we put a ‘0’. Thus a matrix *M* is formed with binary entries indicating the presence and absence of bird species in different districts. We used ‘R ‘, an open-source software (R Core Team, 2021) for the further analysis of the data.

#### 2.3.1. Statistical analysis to identify ecological clusters based on bird diversity

We first identified the similarities between districts in terms of the presence and absence of different bird species. In order to visualize the similarities, we used multidimensional scaling (MDS) (Tao Shen, 2009), a non-linear dimensionality reduction technique. Using MDS on *M*, each district was represented in two dimensions. Subsequently, we applied k-means clustering (MacQueen, 1967) to find out subgroups of districts showing the similarity in the presence and absence of different bird species. We used Euclidean distance as a measure of distance between the districts in their two-dimensional representations for implementing the k-means clustering. The value of *k* has been determined using Elbow method (Thorndike, 1953). To that end, for each value of *k*, we calculated the total within-cluster sum of square (*wss*) considering all the clusters. Subsequently, we plotted *wss* as a function of *k*. We observed that as we increased the value of *k*, the value of *wss* decreased. However, there was significantly less decrease in the value of *wss* after *k* = 4 (see Fig.2). Therefore, we chose *k* = 4 for our clustering. Each of these clusters obtained using the aforementioned method represents one ecoregion.

**Figure 2:**
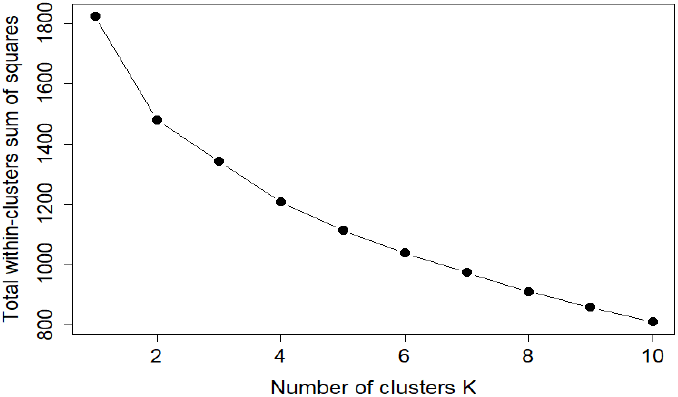
Unsupervised clustering for delineation of ecoregions in Thar desert. The graph represents Within-cluster sum of square (WSS) method for identifying optimal number of clusters. The elbow of the graph was achieved at 4, indicating k in the present k-means analysis would optimally be 4.

#### 2.3.2. Validation of statistically obtained clusters through mathematical analysis

We used the following three different mathematical measures to validate the statistically obtained differences

(a) Cosine similarity matrix: Cosine similarity was calculated between each district using the matrix *M*. Each row in *M* represents the presence/absence data for different bird species for the corresponding district. Hence, we calculated cosine similarities between each pair of rows. This resultant value indicates the similarity between the corresponding districts in terms of the presence/absence of different bird species. If two districts are similar, their cosine similarity value should be close to 1. Otherwise, it should be close to 0.

(b) Map similarity matrix: We also constructed Map similarity matrix that represent the ecoregions with similarity in terms of the presence/absence of different bird species. This matrix is constructed by placing the different districts in rows and columns. If two districts belong to the same ecoregion based on our clustering, the corresponding entry in the matrix is set to ‘1’. Otherwise, the corresponding entry is set to ‘0’.

(c) While the cosine similarity matrix was obtained using matrix *M*, the map similarity matrix was derived based on the results of clustering where in latter we also performed dimensionality reduction. Using both of these techniques, we wanted to find districts having similarities in terms of the presence and absence of bird species. In the map similarity matrix, if two districts were similar, their value was 1, otherwise, the value was 0. In the cosine similarity matrix, if two districts were similar, their cosine similarity value should be close to 1. Otherwise, the cosine similarity value would be close to 0. Therefore, if the cosine similarity matrix and the map similarity matrix perform consistently in identifying similar districts, the difference between an entry in the cosine similarity matrix and the corresponding entry in the map similarity matrix should be close to 0. Hence, we subtracted the entries of the map similarity matrix from the corresponding entries of the cosine similarity matrix. If the entries in the resultant matrix were close to zero, we concluded that the cosine similarity matrix and the map similarity matrix identified the similar and dissimilar districts identically (see Fig.3c).

**Figure 3:**
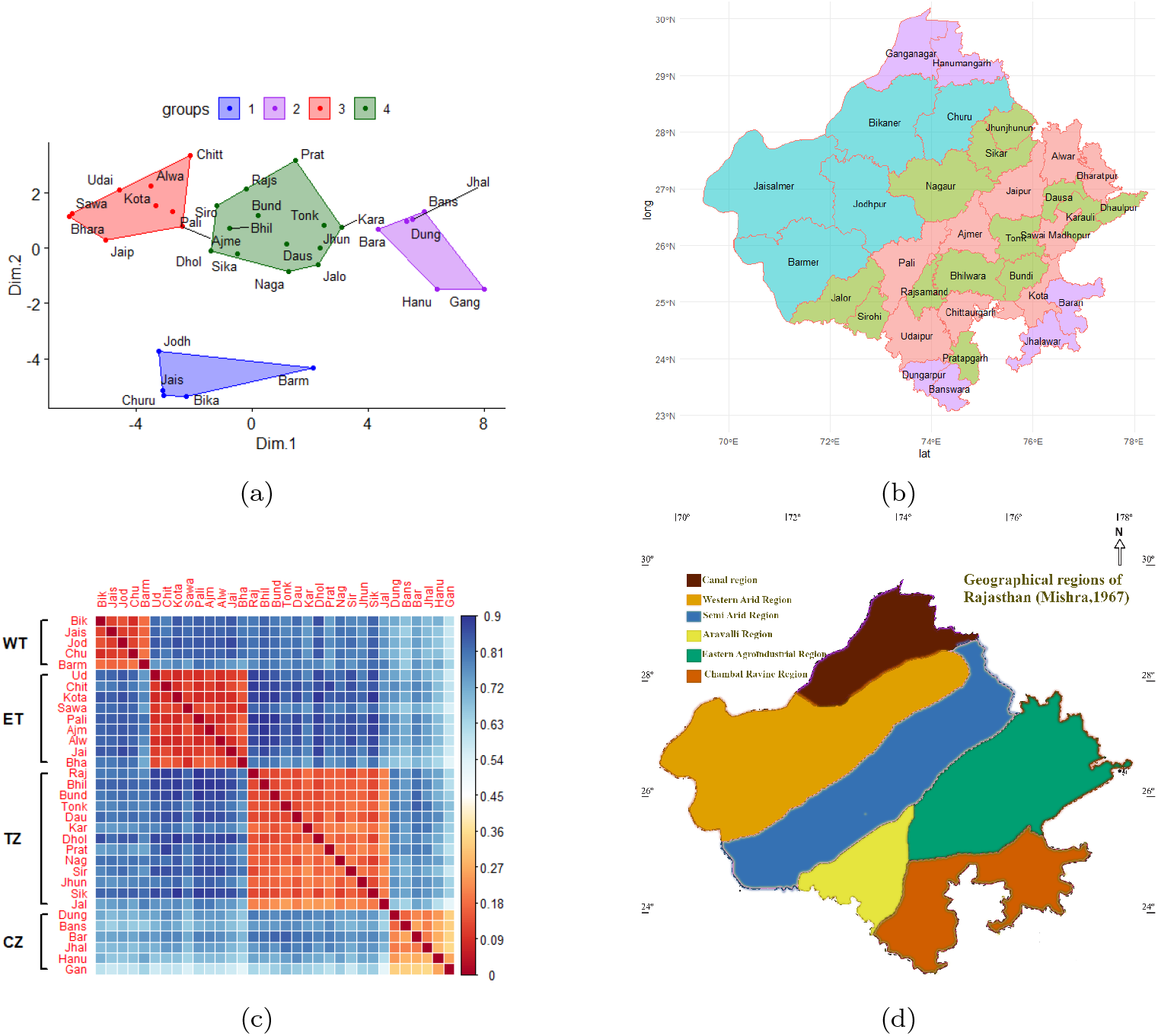
(a) MDS based k-means cluster formed, with *k* = 4, provided four distinct clusters indicated in red, green, purple and blue (b) Each cluster of k-means represents a distinct ecoregion, mapped as blue in Western Thar (WT), pink in Eastern Thar (ET), green in the Transitional Zone (TZ) as depicted in figure 1. The purple is a split zone between two distinct and distant regions formed between south-east Ghagar plains and north west canal region of Thar and is represented as Cultivated Zone (CZ) (see Discussion).(c) Mathematical validation of clusters obtained through unsupervised clustering recapitulates the four clusters. The Figure illustrates a corrplot based on difference between map similarity (0,1) and cosine similarity. The plot scales between 0 to 1, where, values close to 0 (red) indicate districts with very high similarity in avian diversity and those close to 1 (blue) indicate districts that are highly dissimilar.The shade intensity of colours represent extent of similarity or dissimilarity. (d) The 4 ecoregions delineated coincides with four of the seven geographical regions of Thar described earlier. (The map is adapted from Misra (1967))

#### 2.3.3. Spatial diversity

We used a Venn diagram to analyze the spatial diversity of bird species in the ecoregions identified through our clustering method (Fig. 4). Ovals of different colours represent the different ecoregions. While the species shown in the intersections of the ovals represent the common species across the corresponding regions, the species shown in the completely exclusive areas of an oval represent the species unique or exclusive to that ecoregion. Consider *k* number of sets *A*_1_, *A*_2_, …*A*_*k*_. Assume that each set contain the species of one ecoregion. Then the exclusive species of a particular ecoregion *i* is *A*_*i*_ *—* ∪_∀*j≠i*_ (*A*_*i*_ *∩ A*_*j*_). The spatial diversity scales, *α, β*, and *γ* diversity can be analyzed through the Venn diagram. Mathematically, *α* diversity (*N*_*i*_) is the number of species in an individual ecoregion. On the other hand, *γ* diversity (Σ_*i*=*ecoregion*_ *N*_*i*_) is the total number of species in all the ecoregions of Thar and *β* diversity (*ϒ/α*) is the variation of species among the ecoregions.

**Figure 4:**
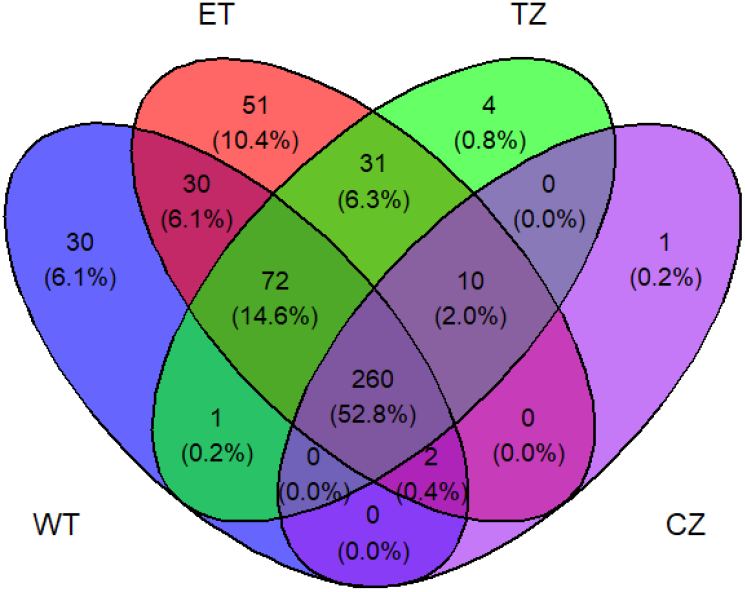
The venn diagram represents the spatial diversity of all the delineated ecoregions. Each oval represents an ecoregion: blue = Western Thar (WT), red= Eastern Thar (ET), green = Transitional Zone (TZ) and purple = Cultivated Zone (CZ). Sum of total counts within all the ovals is the *ϒ* diversity (492). The total number of species under each oval indicates the *α* diversity which are 457 (WT), 395 (ET), 380 (TZ) and 273 (CZ) in the four ecoregions. The ratio of *ϒ* to *α* is the *β* diversity, which is the count difference between the overlapping and unique counts of each ecoregions. For example,it is 1.8 for CZ. Avian diversity common to all the delineated ecoregions is 260. Exclusive species in WT, ET, TZ and CZ were 30, 51, 4 and 1 respectively. Percentage contributions are given in parenthesis.

## 3. Results

The data collected from ebird.org showed presence of 492 species of birds from Thar Desert of Rajasthan across 33 districts. Bharatpur, Sawai Madhopur, Jodhpur and Jaisalmer recorded higher diversity whereas SriGanganagar and Hanumangarh had the least (Table 1). The k-means cluster analysis (Fig. 3a) indicates the presence of four distinct groups 1, 2, 3 and 4 from different districts. Group 1 was formed with nine, Group 2 with thirteen, Group 3 with five and Group 4 with six districts (Table 1). In order to visualize the location of each grouped districts within Thar Desert, the districts from each group were marked within the map of Rajasthan (Fig. 3b). Group 1 mapped to the Eastern Thar (ET), Group 2 districts were located on both sides of the Aravallis forming the Transitional Zone (TZ) and Group 3, was mapped to the Western Thar (WT). Noteworthy, Group 4, was formed of districts located in two extreme ends of the Thar and were geographically not interconnected. We obtained corroborating results of the four groups from both clustering (statistical) as well as mathematical approach of cosine similarity. The difference between the map similarity matrix and cosine similarity matrix is plotted in a corrplot (Fig. 3c). Lesser difference between the two, reflect accuracy of the results. Thus the districts with the highest similarities form separate cluster and indicate the presence of four ecoregions viz. (a) Five districts belong to WT, (b) Nine districts to ET, (c) Thirteen districts to TZ and (d) Six districts to north and southeast of Rajasthan, further named as CZ. All the ecoregions delineated through mathematical analysis were exactly the same as the ones delineated through statistical analysis, thus validating the existence of four distinct ecoregions of Thar. The corrplot (Fig. 3c) also indicates the most closely (red) and distantly (blue) related districts with respect to avian diversity. The deeper the shade, the stronger is the relation. It is evident from the plot that the CZ is comparatively closer to WT and ET than the TZ. Moreover, similarity between the six districts of the CZ is not as strong as amongst districts of other zones. Hanumangarh and Sri Ganganagar show marginal similarity to other four districts (0.24-0.35) of its cluster/group.

**Table 1:** Details of district-wise bird species richness and Multi Dimensional Scaling (MDS) scores in Thar. N is the number of total species, Dimension 1 (Dim.1) and Dimension 2 (Dim.2) are the variables of MDS.

| Zone | Districts | N | Dim.1 | Dim.2 | Zone | Districts | N | Dim.1 | Dim.2 |
| --- | --- | --- | --- | --- | --- | --- | --- | --- | --- |
| Western Thar (WT) | Churu | 328 | -3.06 | -5.34 | Eastern Thar (ET) | Udaipur | 345 | -4.58 | 2.11 |
|  | Bikaner | 311 | -2.3 | -5.4 |  | Sawai Madhopur | 379 | -6.18 | 1.26 |
|  | Jaisalmer | 327 | -3.1 | -5.2 |  | Kota | 312 | -3.33 | 1.53 |
|  | Jodhpur | 323 | -3.2 | -3.7 |  | Bharatpur | 398 | -6.3 | 1.16 |
|  | Barmer | 229 | 2.1 | -4.3 |  | Alwar | 316 | -3.47 | 2.26 |
| Transitional Zone (TZ) | Bhilwara | 275 | -0.77 | 0.72 |  | Pali | 308 | -2.7 | 1.3 |
|  | Bundi | 259 | 0.22 | 1.18 |  | Jaipur | 353 | -5.1 | 0.3 |
|  | Tonk | 229 | 2.48 | 0.83 |  | Ajmer | 303 | -2.4 | 0.8 |
|  | Karauli | 216 | 3.09 | 0.78 |  | Chittorgarh | 298 | -2.1 | 3.4 |
|  | Rajsamand | 272 | -0.22 | 2.14 | Cultivated Zone (CZ) | Dungarpur | 182 | 5.32 | 0.98 |
|  | Nagaur | 241 | 1.27 | -0.86 |  | Hanumangarh | 162 | 6.36 | -1.5 |
|  | Sirohi | 286 | -1.22 | 1.54 |  | Baran | 195 | 4.33 | 0.68 |
|  | Pratapgarh | 245 | 1.5 | 3.2 |  | Sri Ganganagar | 115 | 8.01 | -1.5 |
|  | Jhunjhunu | 222 | 2.3 | 0 |  | Jhalawar | 177 | 5.5 | 1 |
|  | Sikar | 269 | -0.5 | -0.2 |  | Banswara | 167 | 5.9 | 1.3 |
|  | Dholpur | 288 | -1.4 | -0.1 |  |  |  |  |  |
|  | Jalore | 206 | 2.3 | -0.6 |  |  |  |  |  |
|  | Dausa | 249 | 1.2 | 0.2 |  |  |  |  |  |

To determine the factors that could govern formation of these four distinct clusters we next studied the spatial diversity measures within (alpha) and between clusters (beta) of different ecoregions. The venn diagram (Fig. 4) represents the relatedness of each delineated ecoregion with respect to avian diversity. The total avian (*γ*) diversity of Thar as recorded is 492 species with 260 (52.8%) species common to all the ecoregions of Thar. The *α* diversity of ET (457) was highest followed by WT (395), TZ (380) and CZ

(273). Noteworthy, despite the Western Thar being more arid, the *α* diversity was nearly comparable to the TZ and much higher than the Cultivated zone. The *β* diversity that reflects the relative proportion of the total biodiversity was observed to be highest in CZ (1.8) followed by WT (1.29), TZ (1.2) and ET (1.07). The highest number of exclusive species was found in Eastern Thar (10.4%) followed by Western Thar (6.1%), Transitional Zone (0.8%) and Cultivated Zone (0.2%).

Figure 4 also represented the number of exclusive species, i.e., the number of species that exist only in a particular ecoregion and not the others. These species (Table 2) indicate their specific preference to these habitats and thus are indicative of the ecoregion. Surprisingly, despite lower diversity in CZ and TZ, these ecoregions had more presence of endangered and near-threatened exclusive species than WT and ET. The IUCN status of all the exclusive species of WT and ET are recorded as those of ‘least concern’. Amongst all the ecoregions, CZ had the exclusive presence of a near threatened species *Laticilla burnesii* (Rufous-vented grass babbler) whereas TZ was habitat to an endangered species *Athene blewitti* (Forest Owlet) and to *Pelecanus philippensis* (Spot-billed pelican) which is a near threatened species (Table 2).

**Table 2:**
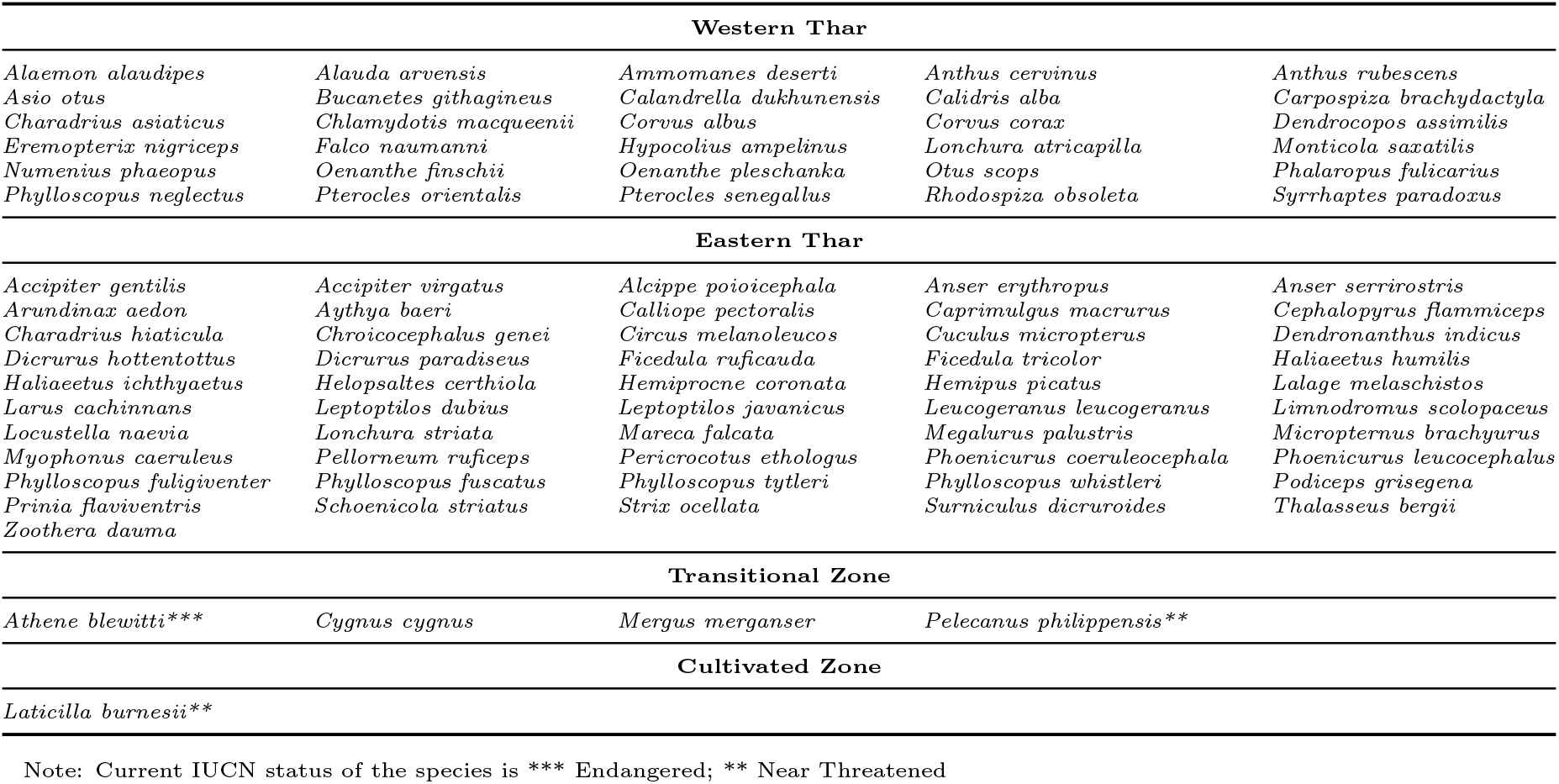
List of bird species specific to each ecoregion in Thar Desert based on the clusters from this study

## 4. Discussion

Ecoregions can be identified by analysing its two major components, the abioitic and biotic. A large number of researchers have relied on climatic and spatial data for delineating ecoregions (Omernik, 1995; Loveland and Merchant, 2004; Kumar et al., 2011). However, for biotic assessments reported so far, ecoregions have been delineated using larger number of taxa including both plants and animals (Bailey, 1983; dos Anjos et al., 2018). Using crowd sourced data from ebird, we report for the first time how a single taxa can be successfully used for delineation of ecoregions. Through a combination of statistical and mathematical methods we demonstrate this approach in defining four ecoregions in Thar which so far was considered as a single ecoregion (Olson et al., 2001). The study highlights (a) birds as a representative biota for inferring shifts in ecoregions (b) the merit of using a continuously updated crowd source information for understanding ecoregion dynamics (c) the effect of anthropogenic activities (d) the need for ecoregion based conservation strategies for protecting both habitat as well as species which are endangered and on the verge of extinction.

Earlier attempts of regionalization in Thar had majorly considered the abiotic components and were aimed primarily to highlight its geographical significance (Misra, 1967; Sharma and Mishra, 2021). Increasing population size (GoR, 2011) and growing urbanization have resulted in greater pace of changes in the abiotic components. While ecoregion maps are considered static, the components governing the ecoregion are conditionally dynamic. Several factors like climate change, anthropogenic activities, etc., can cause dramatic shifts in ecoregions. This indicates possibilities of temporal variability in delineated region, that were not considered in the earlier approaches (Misra, 1967; Sharma and Mishra, 2021). Biotic components due to their pliable and adaptable characteristics are better indicators to capture the spatio-temporal dynamics of ecoregions.

Our study on the 33 districts of Rajasthan that comprise nearly 70% of Thar provided some important insights. Avian diversity based statistical analysis and corresponding mathematical validation from the present study have established the presence of four ecoregions in Thar desert (Fig. 3b and 3c). Earlier methods of k-means clustering using Euclidean distance for delineating geographical ecoregions has used on an extensively large remote sensing data Kumar et al. (2011); Jenerette et al. (2002). Using a comparatively much smaller biota data set with a similar approach we could identify the presence of biotic-ecoregions within the Thar Desert. This was also validated through an orthogonal approach using the cosine similarity method. Using a single taxa on crowd sourced data we could successfully recapitulate four of seven geographical ecoregions (Misra, 1967)) through unsupervised clustering methods. The earlier study was based on assessments of many abiotic characteristics related to physiography, climate, soil, vegetation, agriculture, minerals, population and means of communication. Supervised clustering with *k >* 4 did not resolve the biotic ecoregions into definitive units (Fig. 3d).

Ecoregion WT of the present study corresponds to the Western Arid regions, ET to the Aravalli region and Eastern Agro-Industrial Region and TZ to the Semi Arid Region. The South Eastern agricultural region and Chambal ravine region of Figure 3d have been delineated as the CZ in the present study. This clearly indicates that the variation in bird diversity is a reflection of the differences in the major geographic regions of Thar.

While this remains true for all the other delineated biotic ecoregions of Thar, CZ remains an exception. This ecoregion is composed of districts that are neither geographically interconnected nor similar. The least recorded diversity (*Ni*= 273) and the highest *β* diversity (Fig. 3d and Fig. 4) both indicated a low level of similarity within the districts of this ecoregion. The factors that caused the specific bird diversity and hence formation of this ecoregion could be attributed to the emerging changes of agricultural patterns in Rajasthan. Being the only paddy cultivated Agro-climatic zones of Rajasthan (Sharma, 2013), they attract a large number of specific bird diversity. Agricultural farmlands are known for providing specific habitats to varied bird species. But recent studies have also indicated severe and harmful consequences of intensive farming (Wilson et al., 2005) and overuse of pesticides (Goulson, 2014). Decline in biodiversity of farmlands due to the loss of native habitats (Stanton et al., 2018) can be one possible reason for the formation of CZ ecoregion.

As a vulnerable ecoregion of Thar, CZ is at high risk of habitat fragmentation and raises more concern for inhabiting near threatened species like *Laticilla burnesii* (Table 2). This species is identified as a resident species of this ecoregion and is rapidly decreasing in numbers due to its habitat loss (IUCN, 2022). Some such endangered and near threatened species like *Athene blewitti* and *Pelecanus philippensis* were also identified from TZ that showed specificity to the ecoregion (Table 2). While Aravalli range of ET and TZ have no reports of changing agricultural patterns, it is exposed to another challenge of severe anthropogenic effects. This ecoregion which is known to govern the entire climatic condition of Thar (Roy and Smykatz-Kloss, 2007), is under constant threat of illegal mining and loss of habitat, causing a huge threat to the existence of TZ and its biota.

As Deserts are characterised by relatively fewer species and high endemism, loss of one species is reflected as a much higher percentage of biodiversity loss (Maestre et al., 2021). Thus studies as this become much more important to characterize biota-based ecoregions in desert biomes, identify species specificity and evolve strategies for habitat-specific conservation. The present work has also indicated the importance of biotic-ecoregions in understanding long-term anthropogenic effects and consequences on natural ecosystems. Though the study has a vast scope of globally replicating the approach, there are limitations regarding selection of representative taxa and the requirement of large-scale spatio-temporal biotic data. One could explore the importance of more indicator taxa to understand the variations in ecoregions at different trophic levels and their functions. This framework with different indicator biota is likely to be applicable in any ecosystem. Future studies on variations in spatial diversity scales at different layers of trophic structure shall enable the biota-based sustainability study of a desert ecosystem.

Thar may thus be considered as a desert ecosystem with four distinct ecoregions and needs to be taken into consideration amongst important terrestrial ecoregions. The study also shows birds as an important biota-based tool to understand (a) established ecoregions in an ecosystem, (b)indicator of anthropogenic effects of agri-farmlands and (c) prospects for exploring relation between geographic regions and biota. This work also highlights how a continuous citizens’ engagement could enable generation of large amount of local ecological data for ecoregion specific conservation.

## Acknowledgements

We thank Prof. Santanu Chaudhury, Director, IIT Jodhpur for his support throughout the work. Prof. Surajit Sen from University of Buffalo and and Prof. Anindya Sinha, National Institute of Advanced Studies, for critical review, edits and suggestions. Dr Suman Kundu for data curating. The authors acknowledge funding support to Mitali Mukerji, and financial support to Manasi Mukherjee from the project on Thar-DESIGNS (JCKIF/Thar/Proj-01/2022).

## Authorship contribution statement

**Manasi Mukherjee** contributed in conceptualization, data curation, investigation, methodology, analysis, writing and reviewing. **Angshuman Paul** contributed in methodology, analysis, writing and reviewing and **Mitali Mukerji** contributed in supervision, writing, reviewing and editing.

## References

dos Anjos, L., Volpato, G.H., Lopes, E.V., Willrich, G., Bochio, G.M., Lindsey, B.R.A., Simões, N.R., Mendonça, L.B., Boçon, R., Carvalho, J., Lima, M.R., 2018. Distributions of birds and plants in ecoregions: Implications for the conservation of a neotropical biodiversity hotspot. Austral Ecology 43, 839–849. doi:10.1111/AEC.12626.

Bailey, R.G., 1983. Delineation of ecosystem regions. Environmental Management 7, 365–373. doi:10.1007/BF01866919.

Bharadwaj, G.S., rafi Rahmani, A., 2020. Desert National Park: A Jewel in the Vibrant Thar. Corbett Foundation. URL: https://books.google.co.in/books?id=ld-%5CzgEACAAJ.

eBird, 2023. eBird: An online database of bird distribution and abundance. URL: https://ebird.org/region/IN-RJ?yr=all.

Fraisl, D., Hager, G., Bedessem, B., Gold, M., Hsing, P.Y., Danielsen, F., Hitchcock, C.B., Hulbert, J.M., Piera, J., Spiers, H., Thiel, M., Haklay, M., 2022. Citizen science in environmental and ecological sciences. Nature Reviews Methods Primers 2, 64. URL: https://doi.org/10.1038/s43586-022-00144-4, doi:10.1038/s43586-022-00144-4.

GoR, T.P.D., 2011. Rajasthan at glance. URL: https://urban.rajasthan.gov.in/content/raj/udh/ctp/en/urban-profile/rajasthan-at-glance.html.

Goulson, D., 2014. Pesticides linked to bird declines. Nature 2014 511:7509 511, 295–296. URL: https://www.nature.com/articles/nature13642, doi:10.1038/nature13642.

ul Islam, M.Z., Rahmani, A.R., 2011. Thar Desert, Rajasthan, India: Anthropogenic influence on biodiversity and grasslands. Biodiversity 12, 75–89. doi:10.1080/14888386.2011.585931.

IUCN, 2022. Laticilla burnesii (amended version of 2016 assessment). the iucn red list of threatened species 2022. URL: https://www.iucnredlist.org/species/22735835/111367374.

Jenerette, G.D., Lee, J., Waller, D.W., Carlson, R.E., 2002. Multivariate analysis of the ecoregion delineation for aquatic systems. Environmental Management 29, 67–75. doi:10.1007/s00267-001-0041-z.

Kovařík, P., Pechanec, V., Machar, I., Harmáček, J., Grim, T., 2021. Are birds reliable indicators of most valuable natural areas? Evaluation of special protection areas in the context of habitat protection. Ecological Indicators 132. doi:10.1016/J.ECOLIND.2021.108298.

Kumar, J., Mills, R.T., Hoffman, F.M., Hargrove, W.W., 2011. Parallel k-means clustering for quantitative ecoregion delineation using large data sets, in: Procedia Computer Science, pp. 1602–1611. doi:10.1016/j.procs.2011.04.173.

Lewandowski, E., Caldwell, W., Elmquist, D., Oberhauser, K., 2017. Public perceptions of citizen science. Citizen Science: Theory and Practice 2, 1–3.

Loveland, T.R., Merchant, J.M., 2004. Ecoregions and ecoregionalization: geographical and ecological perspectives. Environmental management 34. doi:10.1007/s00267-003-5181-x.

Lovette, I.J., Fitzpatrick, J.W., 2016. Handbook of Bird Biology. 3 ed., Wiley-Blackwell.

MacQueen, J., 1967. Some methods for classification and analysis of multivariate observations, in: Proceedings of the Fifth Berkeley Symposium on Mathematical Statistics and Probability, University of California Press. pp. 281–298.

Maestre, F.T., Benito, B.M., Berdugo, M., Concostrina-Zubiri, L., Delgado-Baquerizo, M., Eldridge, D.J., Guirado, E., Gross, N., Kéfi, S., Bagousse-Pinguet, Y.L., Ochoa-Hueso, R., Soliveres, S., 2021. Biogeography of global drylands. New Phytologist 231, 540–558. doi:10.1111/NPH.17395.

Misra, V.C., 1967. Geography of Rajasthan, in: Chatterjee, S.P., G., K. (Eds.), India- The land and people. National Book Trust, India, New Delhi, New Delhi. chapter Introducti, p. 183.

Olson, D., Dinerstein, E., Wikramanayake, E., Burgess, N., Powell, G., Underwood, E., D’Amico, J.A., Itoua, I., Strand, H.E., Morrison, J.C., Loucks, C.J., Allnutt, T.F., Ricketts, T.H., Kura, Y., Lamoreux, J.F., Wettengel, W.W., Hedao, P., Kassem, K.R., 2001. Terrestrial Ecoregions of the World. BioScience 51, 933.

Omernik, J.M., 1995. Ecoregions: A Spatial Framework for Environmental Management.

R Core Team, 2021. R: A Language and Environment for Statistical Computing. R Foundation for Statistical Computing. Vienna, Austria. URL: https://www.R-project.org/.

Roy, M.M., Roy, S., 2019. Biodiversity in thar desert and its role in sustainable agriculture. Flora and Fauna) 25, 103–120. doi:10.33451/florafauna.v25i2pp103-120.

Roy, P.D., Smykatz-Kloss, W., 2007. Ree geochemistry of the recent playa sediments from the thar desert, india: An implication to playa sediment provenance. Chemie der Erde 67, 55–68. doi:10.1016/j.chemer.2005.01.006.

Sala, O.E., Chapin, F.S., Armesto, J.J., Berlow, E., Bloomfield, J., Dirzo, R., Huber-Sanwald, E., Huenneke, L.F., Jackson, R.B., Kinzig, A., Leemans, R., Lodge, D.M., Mooney, H.A., Oesterheld, M., Poff, N.L.R., Sykes, M.T., Walker, B.H., Walker, M., Wall, D.H., 2000. Global biodiversity scenarios for the year 2100. Science 287, 1770–1774. doi:10.1126/SCIENCE.287.5459.1770/ASSET/A0209186-7C7A-4372-8261-B95BC35D8854/ASSETS/GRAPHIC/SE0208204003.JPEG.

Sharma, P., 2013. Productivity based allocation of water for irrigation in rajasthan, in: Memoir Geological Society of India. volume 83, pp. 181–196.

Sharma, P.K., Mishra, P., 2021. Geography of Rajasthan. Pareek Publications, Jaipur.

Smits, J.E., Fernie, K.J., 2013. Avian wildlife as sentinels of ecosystem health. Comparative Immunology, Microbiology and Infectious Diseases 36. doi:10.1016/j.cimid.2012.11.007.

Stanton, R.L., Morrissey, C.A., Clark, R.G., 2018. Analysis of trends and agricultural drivers of farmland bird declines in North America: A review. Agriculture, Ecosystems and Environment 254, 244–254. doi:10.1016/j.agee.2017.11.028.

Strimas-Mackey, M., Hochachka, W., Ruiz-Gutierrez, V., Robinson, O., Miller, E., Auer, T., Kelling, S., Fink, D., Johnston., A., 2020. Best practices for using ebird data. URL: https://cornelllabofornithology.github.io/ebird-best-practices/.

Sullivan, B.L., Aycrigg, J.L., Barry, J.H., Bonney, R.E., Bruns, N., Cooper, C.B., Damoulas, T., Dhondt, A.A., Dietterich, T., Farnsworth, A., Fink, D., Fitzpatrick, J.W., Fredericks, T., Gerbracht, J., Gomes, C., Hochachka, W.M., Iliff, M.J., Lagoze, C., Sorte, F.A.L., Merrifield, M., Morris, W., Phillips, T.B., Reynolds, M., Rodewald, A.D., Rosenberg, K.V., Trautmann, N.M., Wiggins, A., Winkler, D.W., Wong, W.K., Wood, C.L., Yu, J., Kelling, S., 2014. The ebird enterprise: An integrated approach to development and application of citizen science. Biological Conservation 169, 31–40. doi:10.1016/J.BIOCON.2013.11.003.

Tao Shen, H., 2009. Multidimensional Scaling, in: cLIU, L., Özsu, M.T. (Eds.), Encyclopedia of Database Systems. Springer US, Boston, MA, p. 1784.

Thorndike, R.L., 1953. Who belongs in the family? Psychometrika 18, 267–276. doi:10.1007/BF02289263.

Wilson, J.D., Whittingham, M.J., Bradbury, R.B., 2005. The management of crop structure: A general approach to reversing the impacts of agricultural intensification on birds? Ibis 147, 453–463. doi:10.1111/j.1474-919x.2005.00440.x.

